# Discovery of A Polymorphic Gene Fusion via Bottom-Up Chimeric RNA Prediction

**DOI:** 10.1101/2023.02.02.526864

**Authors:** Justin Elfman, Lynette Goins, Tessa Heller, Sandeep Singh, Yuh-Hwa Wang, Hui Li

## Abstract

Gene fusions and their chimeric products are typically considered hallmarks of cancer. However, recent studies have found chimeric transcripts in non-cancer tissues and cell lines. In addition, efforts to annotate structural variation at large scale have found examples of gene fusions with potential to produce chimeric transcripts in normal tissues. In this report, we provide a means for targeting population-specific chimeric RNAs to enrich for those generated by gene fusion events. We identify 57 such chimeric RNAs from the GTEx cohort, including *SUZ12P1-CRLF3 and TFG-ADGRG7*, whose distribution we assessed across the populations of the 1000 Genomes Project. We reveal that *SUZ12P1-CRLF3* results from a common complex structural variant in populations with African heritage, and identify its likely mechanism for formation. Additionally, we utilize a large cohort of clinical samples to characterize the *SUZ12P1-CRLF3* chimeric RNA, and find an association between the variant and indications of Neurofibramatosis Type I. We present this gene fusion as a case study for identifying hard-to-find and potentially functional structural variants by selecting for those which produce population-specific fusion transcripts.

**KEY POINTS:**

- Discovery of 57 polymorphic chimeric RNAs
- Characterization of SUZ12P1-CRLF3 polymorphic chimeric RNA and corresponding rearrangement
- Novel bottom-up approach to identify structural variants which produce transcribed gene fusions

## INTRODUCTION

Chimeric RNAs are perhaps most canonically known to arise as products of gene fusion events. Both gene fusions and their associated chimeric RNAs and proteins have been successfully utilized as markers for different cancers and cell types. However, recent studies have also revealed another layer of complexity in that chimeric RNAs can be products of intergenic splicing, in the absence of gene fusion (1, 2). Some of the most impactful findings in this field have stemmed from the discovery of chimeras such as *JJAZ1-JAZF1* (3, 4), *PAX3-FOXO1* (5), and more recently, *EML4-ALK* (6), which mirror known oncogenic gene fusions in the absence of their associated changes at the DNA level. These findings have been further supplemented by read-through chimeras such as *SLC45A3-ELK4* (7) and chimeras such as *ASTN2-PAPPA_AS_* (8), which are produced without noted changes to subject DNA and have demonstrated functions relevant to cancer pathogenesis. There has been a significant effort to catalogue chimeric RNAs in various diseases and cell types to identify such novelties (9–16).

In parallel to our examination of chimeric RNAs which result from splicing phenomena at the RNA level, there is a growing body of research which aims to characterize structural variation genome-wide (17). Structural variant (SV) detection is notably difficult, especially within repetitive regions and in annotation of complex variants and inversions. Some recent efforts have worked to address these issues through parallel examination of a large number of genomes (18) and precise examination of a handful of genomes through integration of short and long-read sequencing (19).

This type of research forms a valuable foundation for downstream analysis, and it is expected that the integration of SV annotation with biological data will provide important steps forward in interpreting the significance of these data. Recent studies have linked structural variation with gene expression in healthy individuals from the GTEx cohort, demonstrating especially strong effects from variants which affect annotated genes (20, 21). While identification of SVs which produce gene fusions are comparatively rare in these studies, these examples tend to be highlighted as findings more likely to produce functional products (22).

There are a number of methods which have aimed to take advantage of transcriptomic data to predict SVs (23–25), and those which specifically leverage chimeric transcript prediction to predict gene fusion events (26–28). Collectively, these methods can be classified as bottom-up approaches, as they identify evidence of the transcript, and seek to trace it back to the change at the DNA level. While bottom-up approaches miss fusions which are not transcribed in the test model, they enrich for gene fusions which affirmatively produce transcripts.

We demonstrate the value of identifying polymorphic SVs with a bottom-up approach by enriching for chimeric RNAs which stratify by population. Not only does this prediction provide value in genotyping and population stratification, but it specifically filters for variants which affect parental genes and produce chimeric RNAs. In this report, we discover *SUZ12P1-CRLF3*, and present it as an example of a polymorphic chimeric RNA. We explore its prevalence across global populations, and provide exemplary characterization of the fusion event and its transcript.

## MATERIAL AND METHODS

### Data Acquisition

RNA-seq and corresponding whole genome sequencing data were obtained from the GTEx project (V6 dbGaP Accesstion phs000424.v6.p1), and the Geuvadis Consortium repository via the 1000 genomes data hub. Expression and sample data for GTEx samples were also obtained from GTEx project (V6 dbGaP Accesstion phs000424.v6.p1).

### Polymorphic ChiRNA predictions

Chimeric RNAs were predicted according to previous publication(9). Briefly, chimeric RNA predictions were generated using the default parameters of EricScript (29), and using GRCh38 as reference. Chimeric RNAs with a low EricScore (<0.6), without predicted breakpoint positions, or with significant (>90% identity) to annotated transcripts were filtered.

In order to enrich for polymorphic chimeric RNAs, we selected chimeric RNAs which were detected in fewer than 250 unique individuals within the GTEx cohort, and within each individual, the chimeric RNA was expressed in more than five unique tissues in more than 66% of samples.

### PCR of Chimeric RNAs

Clinical human leukocytes were obtained in accordance with the Institutional Review Board within the University of Virginia Health system. Whole blood samples were collected from patients admitted to the hospital for non-cancer ailments, and leukocytes were enriched by centrifugation. RNA was extracted using standard protocol for the TRIzol reagent, and reverse transcription was conducted using the SensiFAST kit (Bioline, BIO-65054). qPCR was conducted using the StepOne Plus system (Life Technologies) with SensiFAST SYBR w/ HiRox (Bioline, BIO-92005). Following RT_qPCR and gel electrophoresis, DNA was extracted using the PureLink Gel Extraction Kit (Invitrogen, K210012), and sent to Genewiz for Sanger sequencing.

Patient autopsy samples were obtained from the University of Virginia health system under Institutional Review Board protocol. cDNA was produced via the same methodology detailed above, and PCR was conducted using the standard protocol for Bioline MyTaq™ Red Mix (BIO-25043).

### PCR genotyping of Leukocytes

PCR genotyping was performed utilizing SNPs rs16891982 and rs1426654. rs16891982_G and rs1426654_A were considered to correspond to the European grouping, while rs16891982_C and rs1426654_G considered to correspond to the Non-European grouping. We ordered ThermoFisher Scientific probes C___2842665_10 and C___2908190_10, and performed the assay with the Bioline SensiFAST SYBR Hi-Rox Kit (BIO-92005) per standard protocol. In order to assign a genotype, we required the result to exhibit a standard amplification curve and be consistent across all replicates. Additionally, samples were designated European or Non-European only if the designated genotype for both alleles were in agreement. Otherwise, samples were labelled Unknown.

PCR genotyping targeting the *SUZ12P1-CRLF3* rearrangement was conducted on all samples identified as positive by the TaqMan genotyping assay, as well as 10 samples identified as negative for the rearrangement. We designed four primers to amplify around breakpoints on either side of the inversion (Figures 4D and 4E), and used the standard protocol for Bioline MyTaq™ Red Mix (BIO-25043) for amplification.

Primers used in this study are provided in Supplementary Table S1.

### *In-Silico* AGREP genotyping

The base AGREP function was used to compare a set of query sequences against quality-filtered fastq files. Reads identified to match the query sequences were then aligned with BLAT using parameters minMatch = 1, minScore = 90, minIdentity = 90, and filtered using default parameters of the BLAT suite function repsPSL. These outputs were compared to a 100kb region surrounding the rearrangement. Hits were designated true-positive if they mapped exclusively to the locus, falsepositive if mapped exclusively to other loci, and uncertain if the read mapped to multiple different loci, as long as one was the target locus. Three 70 base pair query sequences were designed to map onto the region spanning breakpoint 1 (SC_Ctrl_), within the deleted region of the breakpoint (SC_Del_), and spanning breakpoints 2 and 3 (SC_Inv_) (Supplementary Figure S1).

### Read alignment and visualization

Reads were quality-filtered using default parameters of the NGSQC toolkit (30), and aligned to GRCh38 with BWA (31). Split and discordant reads were extracted using Samtools (31), and visualized based on pair orientation in IGV (32).

### Secondary structure prediction

We performed predictions centered on breakpoint 1 (chr17:30,770,904), breakpoint 2 (chr17:30,775,728), and breakpoint 3 (chr17:30,778,956) using the methodology described by Szlachta et al(33). Two sets of parameters were considered: predictions within +/- 1000 nucleotides with a 300 nucleotide window and 150 nucleotide step; and +/- 200 nucleotides with a 30 nucleotide window and 1 nucleotide step. Additionally, we performed prediction using both positive and negative strands surrounding rs145766379.

## RESULTS

### The SUZ12P1-CRLF3 RNA transcript is predominantly expressed in populations of African heritage

#### Identification and validation of polymorphic chimeric RNAs

In order to predict polymorphic chimeric RNAs, we utilized 9496 RNA sequencing (RNA-seq) samples from the GTEx cohort (V6), which encompasses 53 different tissues across 549 unique donors. We applied our chimeric prediction pipeline (34), and additionally filtered for chimeras predicted in fewer than 250 individuals, but present in more than five unique tissues in 66% of those expressing the chimera in order to enrich for transcripts found pervasively, but only within particular populations (Figure 1A). These restrictions yielded a total of 57 predictions, including two chimeric RNAs which were stratified by the reported race of donors, *SUZ12-CRLF3* and *TFG-ADGRG7* (Figure 1B). Of these, *SUZ12-CRLF3* showed a statistically significant difference, and is enriched in the donors with African heritage. Though not statistically significant in this study, *TFG-ADGRG7* has been previously reported as a chimeric RNA specific to European ancestry (35).

**Figure 1.**
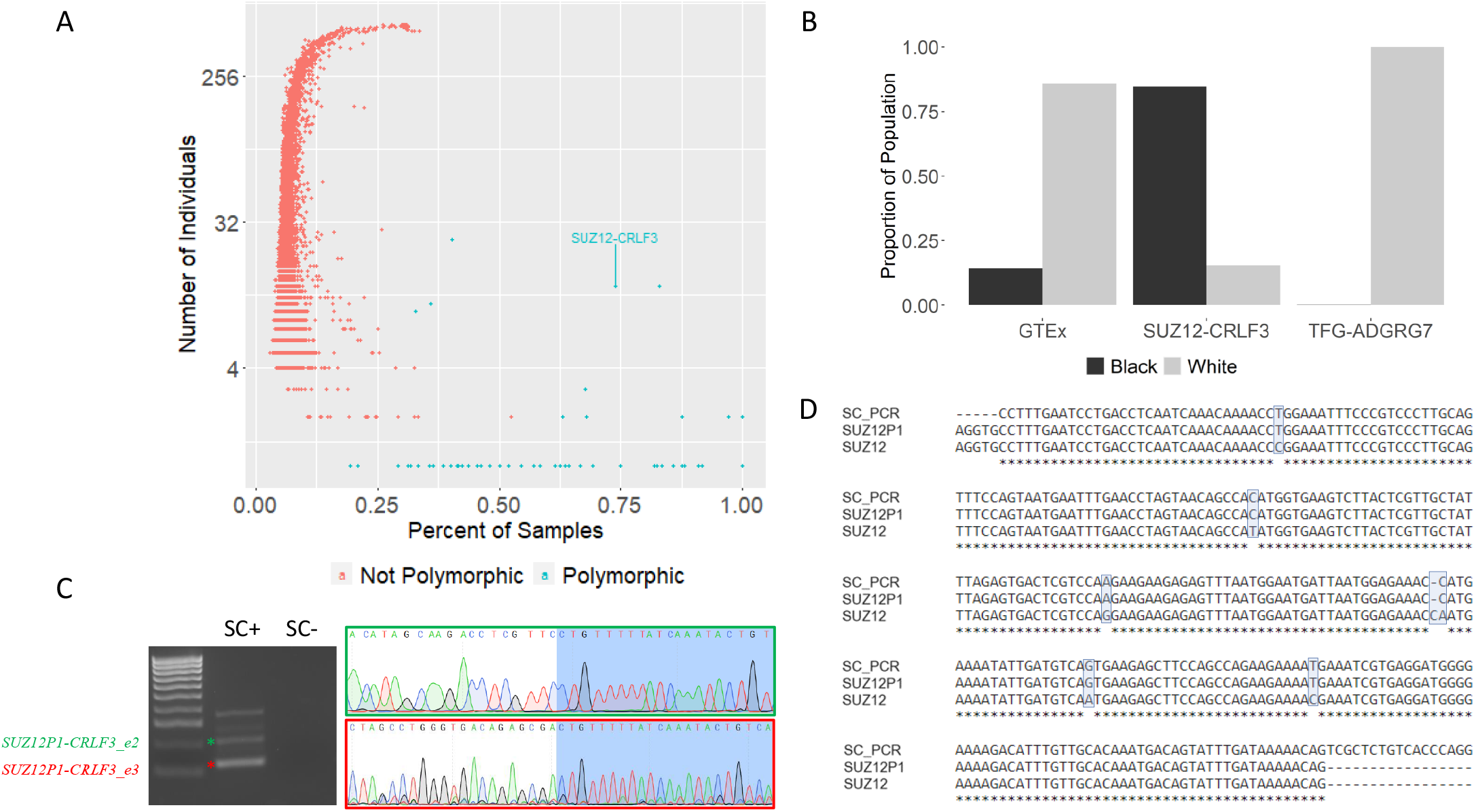
Identification and validation of the SUZ12P1-CRLF3 chimeric RNA. A) GTEx chimeric RNAs by the number of unique individuals in which they are found and the mean proportion of samples for each individual in which they are found. Chimeric RNAs in green have been selected for by the polymorphic filter, those in red have been excluded. B) Racial demographics of the GTEx cohort and subsets of the GTEx cohort including those who express *SUZ12P1-CRLF3* or *TFG-ADGRG7.* C) RT-PCR and Sanger Sequencing supporting the validation of two *SUZ12P1-CRLF3* isoforms (asterisks). D) Transcript sequence confirms the chimeric RNA retains sequence from *SUZ12P1* rather than *SUZ12*.

PCR validation of *SUZ12-CRLF3* revealed a transcript formed not by the *SUZ12* polycomb gene, but instead by its pseudogene *SUZ12P1* and *CRLF3.* We identified two isoforms of this transcript, with junction sequences comprised by the joining of *SUZ12P1* exon 8 to *CRLF3* exon 2 or 3 (Figure 1C and D).

We sought to validate these results in populations from the 1000 genomes cohort which had RNA-seq in a subset of samples. We directly compared 28 base pairs spanning the junction sequence of each isoform of the predicted *SUZ12P1-CRLF3* chimera to quality-filtered RNA-seq reads. We found *SUZ12P1-CRLF3* in the Nigerian population (YRI) at a similar rate to what was found in African-American GTEx samples, and was not found in any of the four European populations (Table 1).

**Table 1.** SUZ12P1-CRLF3 occurrence in 1000 genomes populations.

| ***Cohort*** | ***Description*** | ***Detected*** | ***Total*** | ***%*** |
| --- | --- | --- | --- | --- |
| *CEU* | *Northern European (Utah)* | *0* | *92* | *0* |
| *FIN* | *Finnish in Finland* | *0* | *95* | *0* |
| *GBR* | *British in England and Scotland* | *0* | *95* | *0* |
| *TSI* | *Toscani (Italy)* | *0* | *93* | *0* |
| ***YRI*** | ***Yoruba in Ibadan (Nigeria)*** | ***13*** | ***89*** | ***14.6*** |
| *GTE<sub>x</sub>* | *White* | *2* | *462* | *0.4* |
| ***GTE<sub>x</sub>*** | ***Black or African-American*** | ***11*** | ***77*** | ***14.3*** |
| *GTE<sub>x</sub>* | *Other/Unknown* | *0* | *10* | ***0*** |

#### Validation and enumeration of the SUZ12P1-CRLF3 chimeric RNA in clinical blood samples

We expanded our validation to a set of clinical leukocyte samples collected by the University of Virginia Health System from patients admitted for non-cancer ailments. This dataset consists of 1351 total samples, comprising 948 unique donors, 219 of whom have provided samples at more than one time point (Figure 2A). Reflecting patient population this cohort is predominantly male and white (Figures 2B and 2D), and the mean age of male donors significantly exceeds the mean age of female donors (Figure 2C). Additionally, clinical codes are reported for these donors, which allows for visibility into clinical phenotypes for donors. We find that the median code is listed in 24 unique instances and the 24 most common codes accounting for approximately half of all annotated codes (Figure 2E), and the median donor possesses six unique codes (Figure 2F).

**Figure 2.**
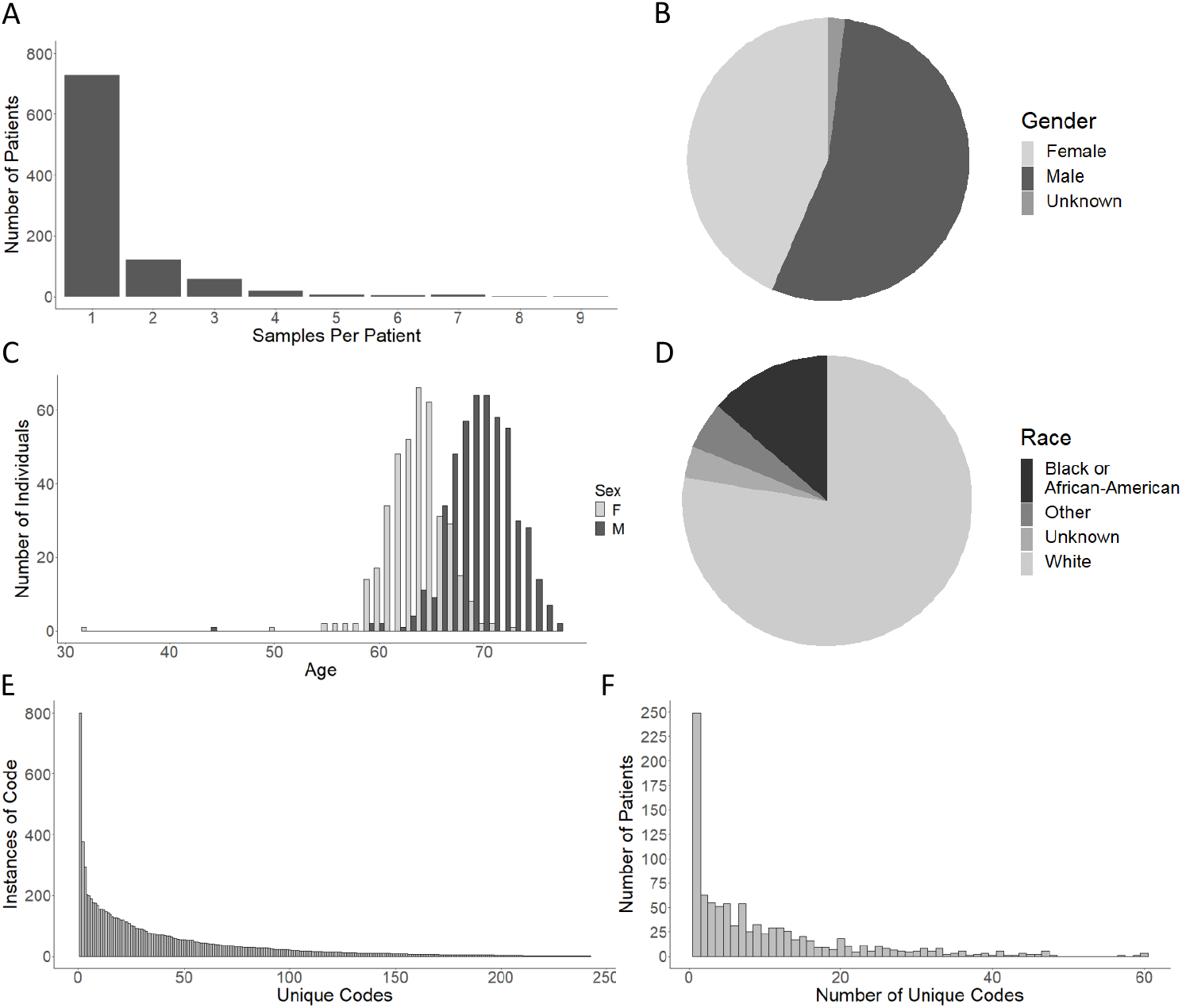
1351 clinical donor leukocytes collected from the UVA Health System. A) The number of samples per unique donor. B) Gender distribution of the dataset. C) Age distribution of the dataset, by listed sex. D) Racial distribution of donors, as annotated. E) The number of recurrences per unique clinical code. F) The total number of clinical codes exhibited per patient.

We designed a TaqMan qPCR assay to detect both isoforms of *SUZ12P1-CRLF3* (Figure 3A), and found that *SUZ12P1-CRLF3* was expressed significantly more often in patients with a reported race of black or African American than in patients reported as white or other (Figure 3C). However, the proportion of individuals categorized as black or African-American were considerably lower than observed in the GTEx and 1000 genomes cohorts.

**Figure 3.**
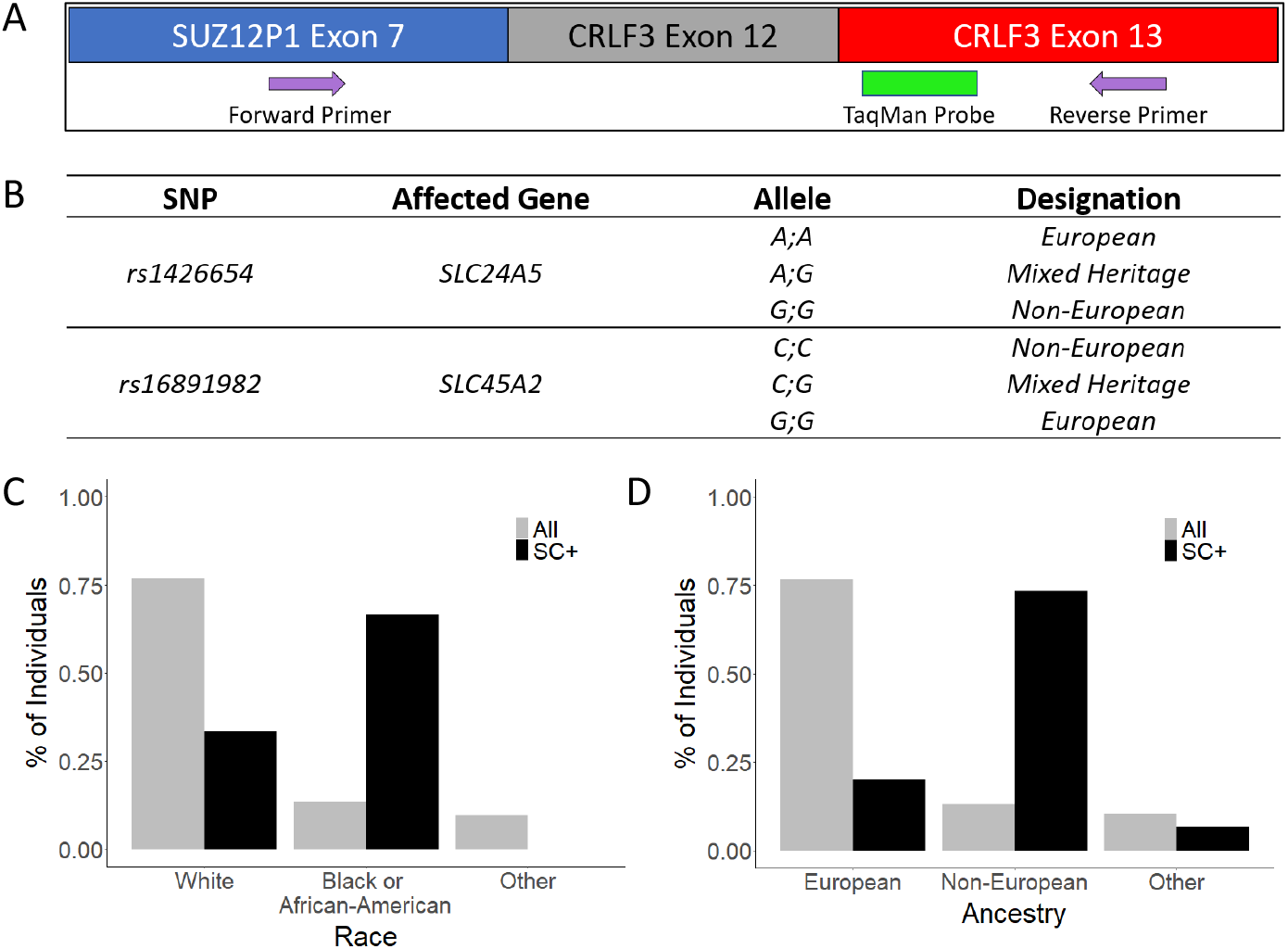
Characterization of SUZ12P1-CRLF3 in clinical samples. A) Probe-based qPCR design. B) SNPs used to genotype leukocyte donors paired with their heritage designation. C) Proportion of population expressing the *SUZ12P1-CRLF3* chimeric RNA compared to the proportion of the population classified by annotated race prior to SNP genotyping and D) by annotated heritage following SNP genotyping.

Noting that clinical racial classifiers are not indicative of ancestry, we sought to more accurately stratify these groupings by ancestry informative markers, rs16891982 and rs1426654, which correlate strongly with population (36) (Figure 3B). We utilized TaqMan SNP Genotyping Assay probes to test all samples expressing *SUZ12P1-CRLF3*, as well as a panel of both white and African-American individuals not expressing *SUZ12P1-CRLF3.* In all cases where sample genotyping was successful, the population designations for rs1426654 and rs16891982 were in agreement, and of these, the sample designation was changed in six cases. In five of these cases, a designation of unknown race was changed to non-European, while in one case, a designation of white was changed to non-European (Supplementary Table S2). After SNP genotyping correction, we found that donors expressing *SUZ12P1-CRLF3* were more likely to possess genetic heritage from non-European populations, approaching a similar proportion (10.1%) to that observed in the GTEx African-American and 1000 genomes YRI populations (~14.5%) (Figure 3D).

### The *SUZ12P1-CRLF3* RNA is generated by a genomic rearrangement at 17q11.2

#### The SUZ12P1-CRLF3 transcript is found in multiple tissues of a SUZ12P1-CRLF3+ Donor

In GTEx data, the *SUZ12P1-CRLF3* transcript was found in multiple tissues of each expressing donor. We sought to confirm this observation experimentally. We obtained autopsy samples from 13 unique black donors within the University of Virginia Health System, and found that one individual expressed *SUZ12P1-CRLF3.* We then obtained samples from 11 different tissues from this *SUZ12P1-CRLF3+* donor, and found the RNA expressed ubiquitously across tissues (Figure 4A).

**Figure 4.**
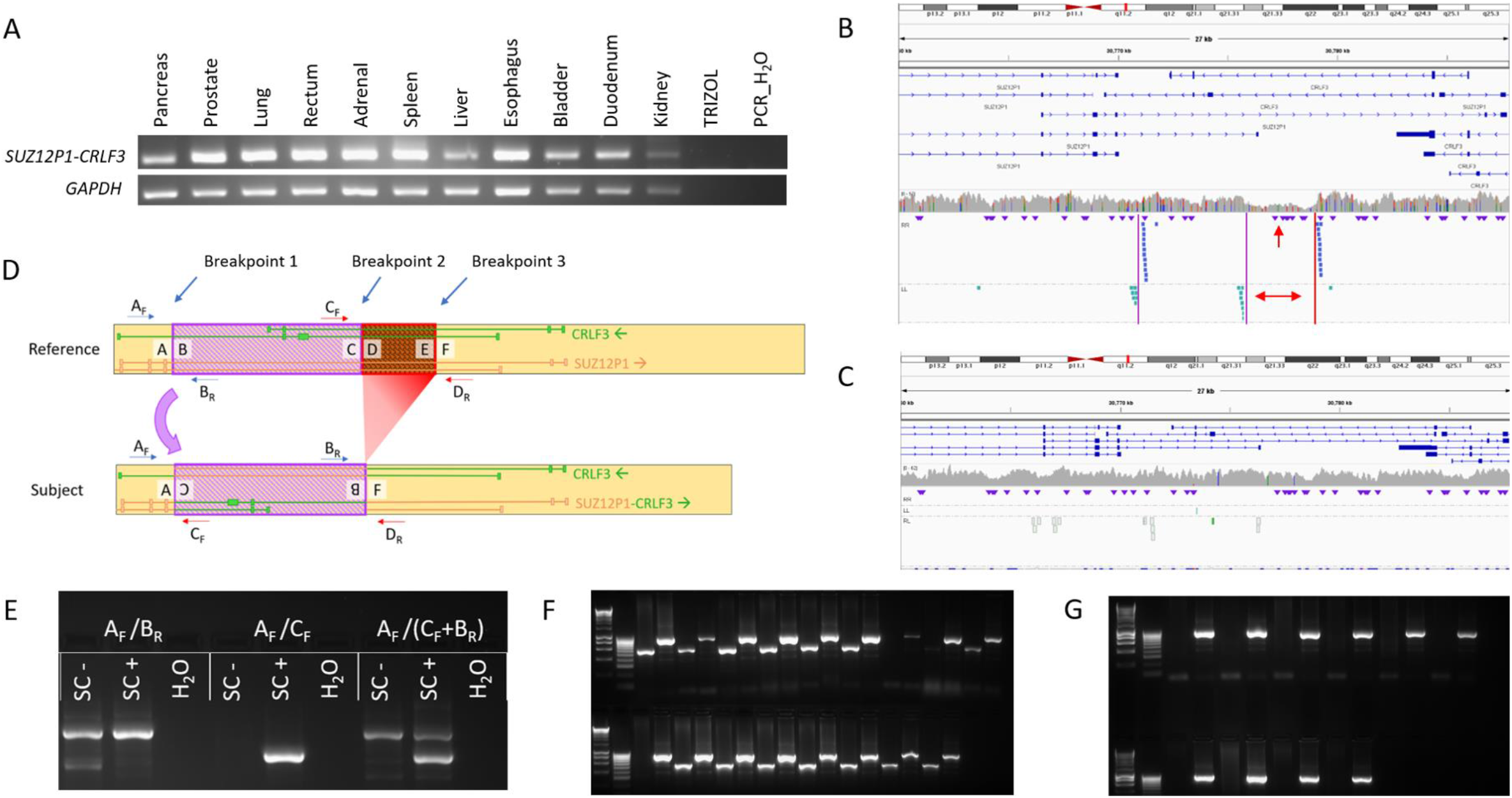
Characterization of the SUZ12P1-CRLF3 chromosomal rearrangement. A) The *SUZ12P1-CRLF3* chimeric RNA is detected in all provided tissues of a single donor. B) Discordant read alignment reveals an inversion and adjacent deletion at the *SUZ12P1-CRLF3* locus in an individual expressing the chimeric RNA, and C) are absent in an individual who does not express the chimeric RNA. D) Schematic of the *SUZ12P1-CRLF3* chromosomal rearrangement, representation of the chimeric RNA, and primer design for genotyping PCR. E) Genotyping PCR. F) Genotyping assay applied to 17 *SUZ12P1-CRLF3-expressing* samples. G) Genotyping assay applied to 10 non-*SUZ12P1-CRLF3*-expressing samples.

#### Discordant read alignment reveals a complex rearrangement at the SUZ12P1-CRLF3 locus

SUZ12P1 and CRLF3 are located in opposing orientations on the same locus within 17q11.2, and several isoforms overlap coordinates. This proximity, paired with its consistent rate of detection, population specificity, and lack of tissue specificity formed a strong basis that the *SUZ12P1-CRLF3* chimeric RNA may be generated by a polymorphic rearrangement. In order to confirm this, we extracted split and discordant reads from all 13 GTEx samples which express *SUZ12P1-CRLF3* as well as 11 random GTEx samples which do not express *SUZ12P1-CRLF3.* Alignment of these reads to HG38 revealed a pile-up within a region on 17q11.2, overlapping isoforms of both *SUZ12P1* and *CRLF3*. The alignment of the left-left and right-right reads indicate an inversion, and the spacing of the right-right reads around the downstream inversion breakpoint, as well as the depletion of reads aligning to this region indicates a deletion in the subject genome (Figures 4B and 4C).

#### Genotyping for the SUZ12P1-CRLF3 Rearrangement

We developed a PCR genotyping assay to amplify over each breakpoint for both the reference and subject genomes (Figures 4D and 4E), which utilized the exchange in the orientation of primers B_R_ and C_F_ caused by the inversion. This assay was performed on the *SUZ12P1-CRLF3+* panel as well as the panel of white samples not expressing *SUZ12P1-CRLF3.* Genotyping revealed that all but two individuals expressing the *SUZ12P1-CRLF3* chimeric RNA were heterozygous for the alternate allele whereas all *SUZ12P1-CRLF3* negative cases had no amplification for the alternate allele (Figures 4F and 4G).

#### Bioinformatic Evaluation of the SUZ12P1-CRLF3 Rearrangement

Next, we developed a pipeline to perform *in-silico* genotyping on whole genome sequencing data by string-matching unique genomic sequence to identify supporting reads, and mapping those reads back to the reference genome (Supplementary Figure S2). We applied the *in-silico* genotyping pipeline to the 1000 genomes cohort (Figure 5A) using these sequences. As a control, we also queried a sequence spanning the DNA junction of the *TFG-ADGRG7* fusion, and found that these variants are invariably detected in European and admixed populations, consistent with previous observation (35). In contrast, the *SUZ12P1-CRLF3* variant was found exclusively in the African (AFR) superpopulation, including ACB, ASW, ESN, GWD, LWK, MSL, and YRI subpopulations, with the exception of one individual in the admixed MXL population (5B and 5C). Additionally, all AFR subpopulations exhibited minor allele frequencies > 5%, with allelic distribution within Hardy-Weinberg equilibrium (Supplementary Table S3).

**Figure 5.**
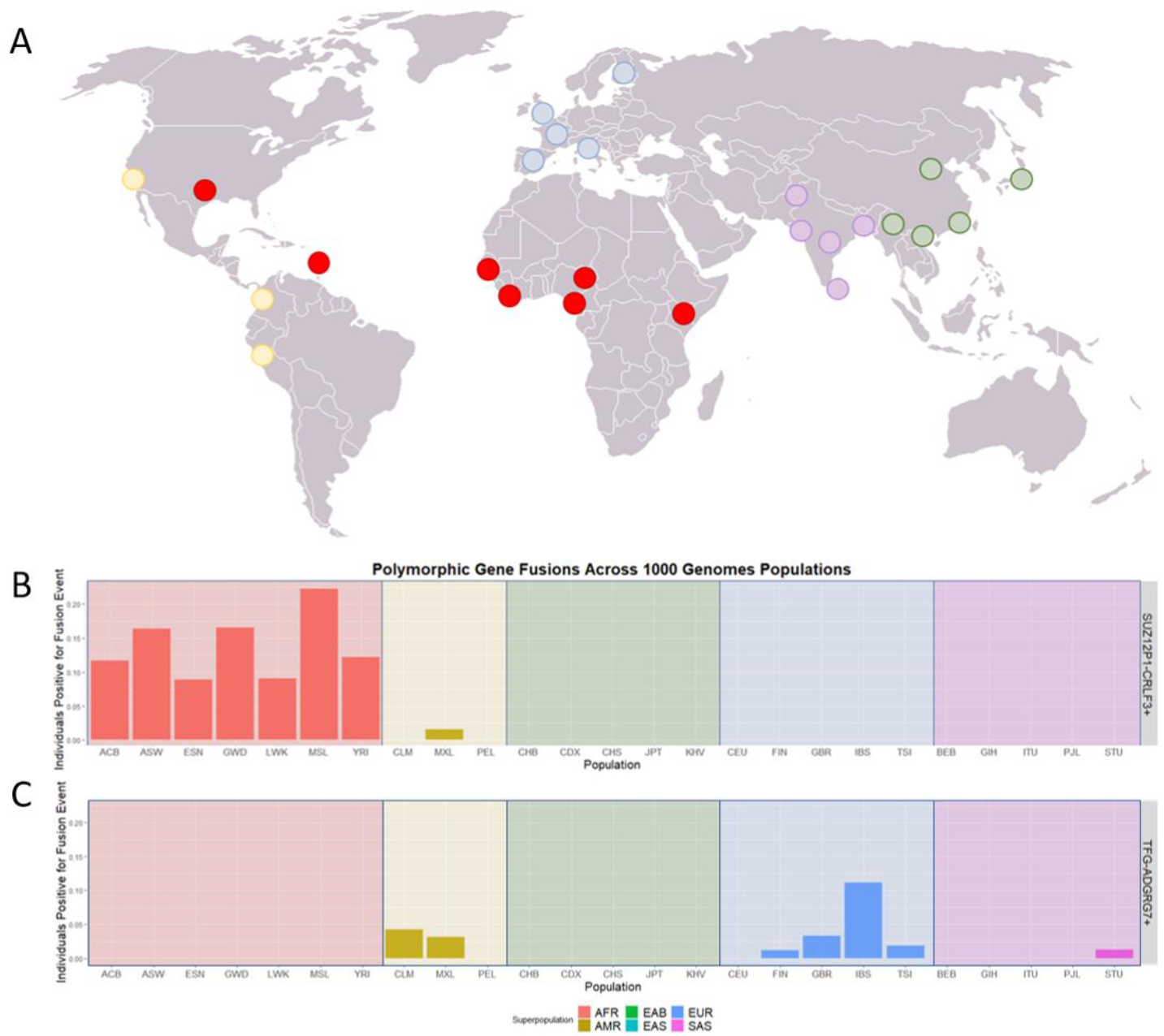
In silico genotyping of the SUZ12P1-CRLF3 variant in 1000 genomes cohort. A) 1000 genomes cohort populations, sorted by superpopulation. B) Frequency of variant allele detection by subpopulation and superpopulation for *SUZ12P1-CRLF3* and C) *TFG-ADGRG7.*

### Functional implications of the *SUZ12P1-CRLF3* rearrangement

#### Absence of association between SUZ12P1-CRLF3 allele and gene expression

We found no difference between individuals with the rearrangement allele in height, weight, BMI, or age in neither GTEx nor leukocyte donor cohorts after accommodating for differences in these measurements by gender (Supplementary Figure S3). In the GTEx population, we did not find any difference in expression in *SUZ12P1* or *CRLF3* expression (Figure 6A and Figure 6B) nor any clear difference in exon usage in terms of splicing isoforms between *SUZ12P1-CRLF3+* and *SUZ12P1-CRLF3-* groups within an African-American background (Supplementary Figure S4). Additionally, we found no significant difference in expression of other genes within this locus (Supplementary Figure S5).

**Figure 6.**
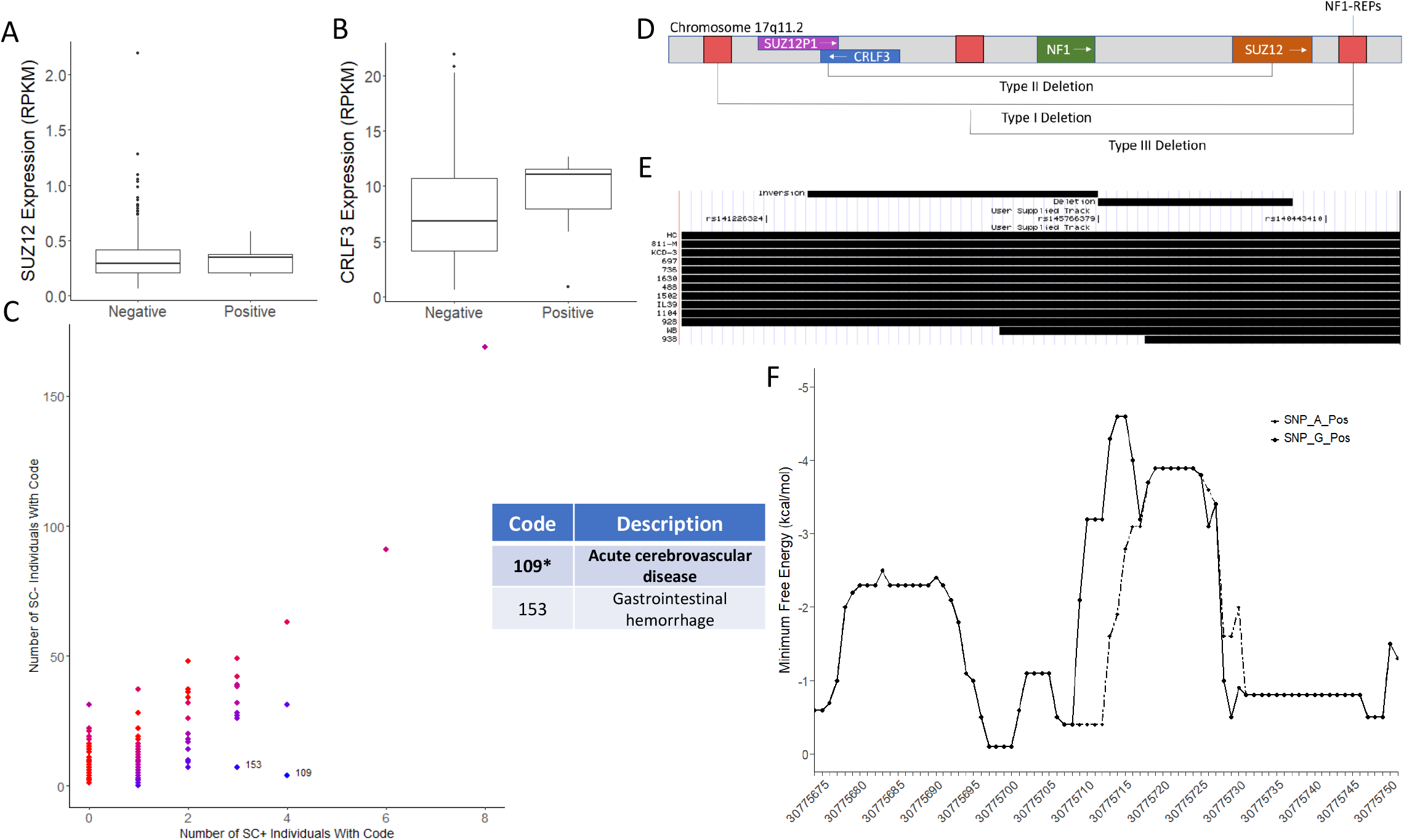
Functional correlations of the SUZ12P1-CRLF3 variant. A) *SUZ12P1* and B) *CRLF3* expression by rearrangement allele. C) Clinical codes associated with leukocyte donors. *FDR-adjusted p-value = 0.049. D) Annotation of genes and repetitive elements on 17q11.2 relevant to microdeletions in type I neurofibromatosis. E) Microdeletions characterized by Steinmann et al. (2007) overlaid with the *SUZ12P1-CRLF3* variant and associated SNPs at the locus. F) RNAfold secondary structure prediction minimum free energy (MFE) calculations for the locus. Predictions are made using a 30 bp sliding window at each base pair surrounding Rs145766379_A_G and are calculated using reference (A) and variant (G) alleles.

#### Association of SUZ12P1-CRLF3 allele with patient clinical codes

The human leukocyte samples we used have 234 unique clinical codes which describe historical health issues that each patient has exhibited. Of the leukocyte samples tested for the *SUZ12P1-CRLF3* chimeric RNA, 123 samples from African-American individuals had associated clinical codes, including 14 of 15 samples which tested positive for the *SUZ12P1-CRLF3* chimeric RNA. We discovered a strong association of the rearrangement allele with acute cerebrovascular disease (ACD), as well as a weaker association with gastrointestinal haemorrhage (Figure 6C).

Notably, the 17q11.2 locus is associated with Type I Neurofibramatosis (NF1), caused by alterations within or microdeletions which span the *NF1* gene. Vasculopathy is a known indication of NF1, and both cerebrovascular (37–40) and gastrointestinal manifestations (41–44) have been associated with the disease. The three most common deletions occur between repetitive regions (NF1-REPs) surrounding the *NF1* gene (Figure 6D). In contrast, type II deletions occur via non-allelic homologous recombination between regions of the *SUZ12* polycomb gene and the *SUZ12P1* pseudogene. While data covering breakpoints of patients with type II deletions are rare, we found that 15.4% of patients with type II deletions as published previously by Steinmann et al. (45) possessed rearrangements with breakpoints within the coordinates of the SUZ12P1-CRLF3 rearrangement (Figure 6E).

#### Rs145766379 may inform initial cause of rearrangement

Due to scarcity of data characterizing type II NF1 deletions, we searched for a panel of single nucleotide polymorphisms (SNPs) associated with the variant to use as a proxy for the genotype. Significantly correlated SNPs clustered around the immediate locus at 17q11.2 (Supplementary Figure S6). No SNPs associated with the rearrangement were previously associated with disease within ClinVar (46) or Gnomad (47). Three of these SNPs were located in close vicinity to the rearrangement, including Rs145766379_A_G, which is located within the downstream breakpoint region of the inversion (Figure 6E).

Genomic regions with potential to form high-stability secondary structures are highly correlated with double-stranded breaks, especially those mediated by topoisomerase II (33). Consequently, these regions overlap with recurrent rearrangement breakpoints in cancer (48, 49). Therefore, we simulated likely secondary structure formation across this region and found that Rs145766379_A_G introduces a region of increased likelihood for secondary structure formation near the observed breakpoint (Figure 6F).

Additionally, Rs145766379_A_G extends a region of 14 base pair consecutive homology to 19 base pairs (Supplementary Figure S7), perhaps increasing the likelihood for an inversion to occur. This region of increased homology resides within one of several Alu-family repeat elements with overlapping homology, which we believe may have initially mediated the rearrangement. Mapping of split reads reveals a discontinuity of the deleted region, indicating that the inversion precedes the deletion, and we find that the inversion moves an AluSg4 element into alignment with an upstream AluSg, providing a potential means for Alu/Alu recombination, resulting in the final complex rearrangement (Supplementary Figure S7).

## DISCUSSION

Chimeric RNAs have previously been characterized within a number of diseases, with particular emphasis on cancer. Some recent efforts have performed a similar characterization in healthy cohorts in order to create an original resource for chimeric RNA characterization in non-diseased donors (50). We utilized the healthy subgroup to enrich for population-specific chimeras which we could treat as polymorphisms or structural variant calls in correlational analyses. To our knowledge, this is the first analysis of the sort.

In this study, we identified 57 such population-specific chimeric RNAs, including two whose distribution we assessed across the populations of the 1000 genomes cohort. Of these, we performed an exemplary analysis of *SUZ12P1-CRLF3*, including characterization of the rearrangement which produces the chiRNA, and potential mechanisms for its genesis. Interestingly, we identified two donors who tested positive for the chimeric RNA but did not possess the variant genotype, which may suggest that the transcript can be created via intergenic splicing as well as canonical transcription from a fusion gene. This data places *SUZ12P1-CRLF3* within a small group of chimeric RNAs that have been detected in this manner, including *JAZF1-JJAZ1* (3), *PAX3-FOXO1* (5), and *EWSR1-FLI1* (51). Additionally, we performed correlational studies to identify a panel of associated variants and were able to examine its association with ACD and gastrointestinal hemorrhage, forming a basis from which its relevance in NF1 can be further studied.

Our method provides a targeted, bottom-up means for identifying gene fusion transcripts which can be analyzed similarly. This stands in contrast to top-down approaches focusing on variants that overlap gene annotations, which cannot be assumed to produce a fusion transcript, and are potentially limited by variables such as the reference used or blackout regions from alignment. We believe that this methodology will prove valuable when applied to other similar datasets.

One common mischaracterization which plagues this field is the assumption that chimeric RNAs and gene fusions are exclusively markers of cancer or disease. While often true, previous discoveries of chimeric RNAs in normal tissues have carved out exceptions to the rule (5, 52, 53). Similarly, this study provides an example of a fusion transcript produced within healthy populations at rates in accordance with Hardy-Weinberg equilibrium. Each of these chimeric RNAs, therefore, do not seem to provide selective pressure, despite noted associations of *SUZ12P1-CRLF3* with ACD and gastrointestinal haemorrhage. It is possible that there was positive selection for this allele at some point in history that no longer exists.

This report provides a framework for similar characterization studies including the 55 population-specific chimeric RNAs not fully characterized in this study as well as in other model systems and cohorts. We believe that this line of analysis is especially useful as a supplement to topdown structural variant prediction, especially when targeting potentially functional variants which produce fusion genes.

## DATA AVAILABILITY

No new data were generated in support of this research.

## SUPPLEMENTARY DATA

Supplementary Data are available at NAR online.

## ACKNOWLEDGEMENT

The authors would like to acknowledge the BTRF for providing tissue samples to support this work.

