## Supplementary tables and figures for "Discovery of A Polymorphic Gene Fusion via Bottom-Up Chimeric RNA Prediction"

**SUPPLEMENTARY TABLE AND FIGURES LEGENDS**

Supplementary Table 1. *List of Primers Used*.

Supplementary Table 2. *Changes to sample designation resulting from SNP genotying.*

Supplementary Table 3. *Minor allele frequency in 1000 genomes populations with SUZ12P1-CRLF3-expressing individuals.*

Supplementary Figure 1: *AGREP queries and representation on locus cartoon.*

Supplementary Figure 2: *AGREP pipeline diagram.*

Supplementary Figure 3: *Donor characteristics in GTEx and UVA cohorts stratified by SC genotype.* GTEx donor A) height, B) weight, and C) BMI by SC genotype. UVA donor A) height, B) weight, and C) BMI by SC genotype.

Supplementary Figure 4: *SUZ12P1 and CRLF3 read coverage by SC genotype.*

Supplementary Figure 5: *Gene expression of genes on the 17q11.2 locus by SC genotype.*

Supplementary Figure 6: *Manhattan plots of SNPs associated with the SC variant.* A) Displaying variants across HG38, B) on chromosome 17, and C) at the SC locus on 17q11.2.

Supplementary Figure 7: *The SUZ12P1-CRLF3 variant was likely mediated by sequence homology of Alu elements.* A) The *SUZ12P1-CRLF3* locus with annotated Alu elements. The rearrangement was likely caused by an inversion followed by a deletion mediated by B) homologous sequence at each breakpoint, shown in light blue for the inversion and purple for the deletion. Regions containing the possible exact breakpoints are highlighted in green. C) RNAfold predicted secondary structure for the 30 bp window surrounding peak MFE for the reference allele and D) for the variant allele.

Supplementary Table S1

| Primer Name | Sequence |
| --- | --- |
| SUZ12P1-CRLF3_F | GGATGGGGAAAAGACATTTGTTGC |
| SUZ12P1-CRLF3_R | GGGAGTCCAAGGTGCGAAGACTTT |
| SUZ12P1-CRLF3_Taq | GGAACGAGGTCTTGCTATGTTGTCCAAACTG |
| SUZ12P1-CRLF3_$A_F$ | GCCGTACGTAAATCTTGGGG |
| SUZ12P1-CRLF3_$B_R$ | CGTCTGTGTTTCTTGCCATCC |
| SUZ12P1-CRLF3_$C_F$ | TTCCCTAAGACAAGCATTGTAACAC |
| SUZ12P1-CRLF3_$D_R$ | CTCCACCTCTGGGTGACAGA |
| GAPDH_F | CTGACTTCAACAGCGACACC |
| GAPDH_R | TTACTCCTTGGAGGCCATGT |

Supplementary Table S2

| SNP Agreement | | Sample Designation | |
|---|---|---|---|
| *Yes* | *No* | *Unchanged* | *Changed* |
| *15* | *0* | *9* | *6* |

| Changes in Sample Designation | | | | | |
|---|---|---|---|---|---|
| *W → N* | *W → U* | *B → E* | *B → U* | *U → E* | *U → N* |
| *1* | *0* | *0* | *0* | *0* | *5* |

Supplementary Figure S1

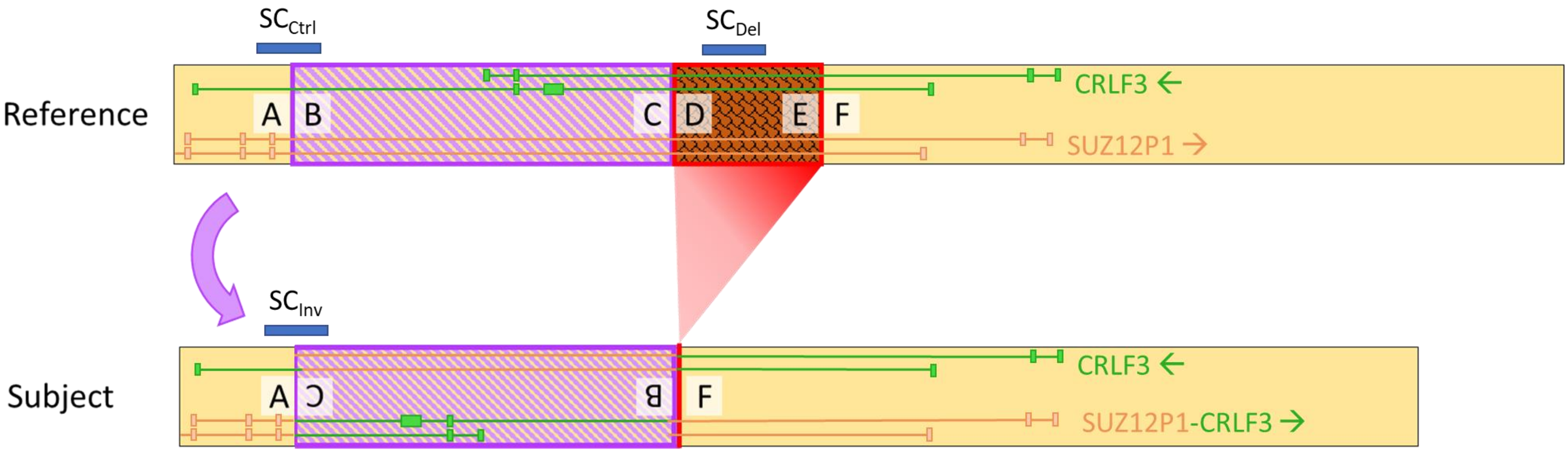


| Query Name | Sequence |
|---|---|
| $SC_{Ctrl}$ | CCTCGCGATCCGCCTGCCTTGGCCTCCCAAAGTGCTGGGATTGCAGGCGTGAGCCACCATGCCTAGGCGA |
| $SC_{Inv}$ | GTGAGCCAAGATCACACCACTGCACTCCATCCTGGGTGACAGAGCAAAAG |
| $SC_{Del}$ | GTTAATATATTTAATATATTGCAGTGAGGA |

Supplementary Table S3

| Population | SC/SC | SC/Ref | Ref/Ref | Total | MAF |
|---|---|---|---|---|---|
| ACB | 3 | 8 | 83 | 94 | 7.4% |
| ASW | 2 | 7 | 46 | 55 | 10.0% |
| ESN | 2 | 5 | 72 | 79 | 5.7% |
| GWD | 4 | 14 | 91 | 109 | 10.1% |
| LWK | 2 | 7 | 90 | 99 | 5.5% |
| MSL | 2 | 4 | 21 | 27 | 14.8% |
| MXL | 0 | 1 | 63 | 64 | 0.7% |
| YRI | 2 | 8 | 72 | 82 | 8.3% |

Supplementary Figure S2

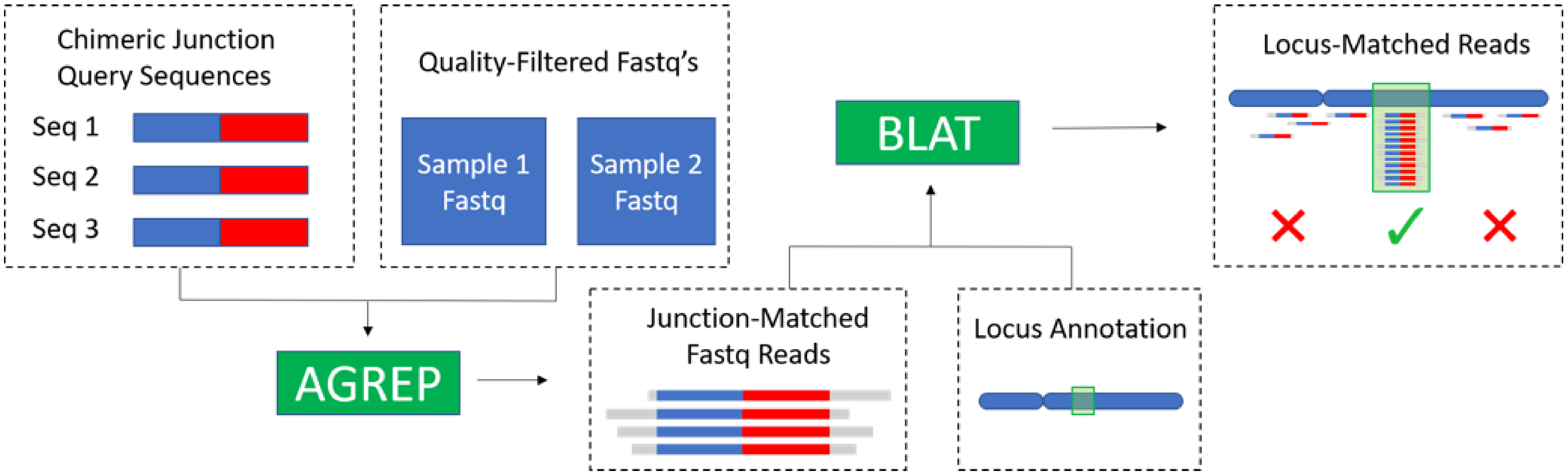

Supplementary Figure S3

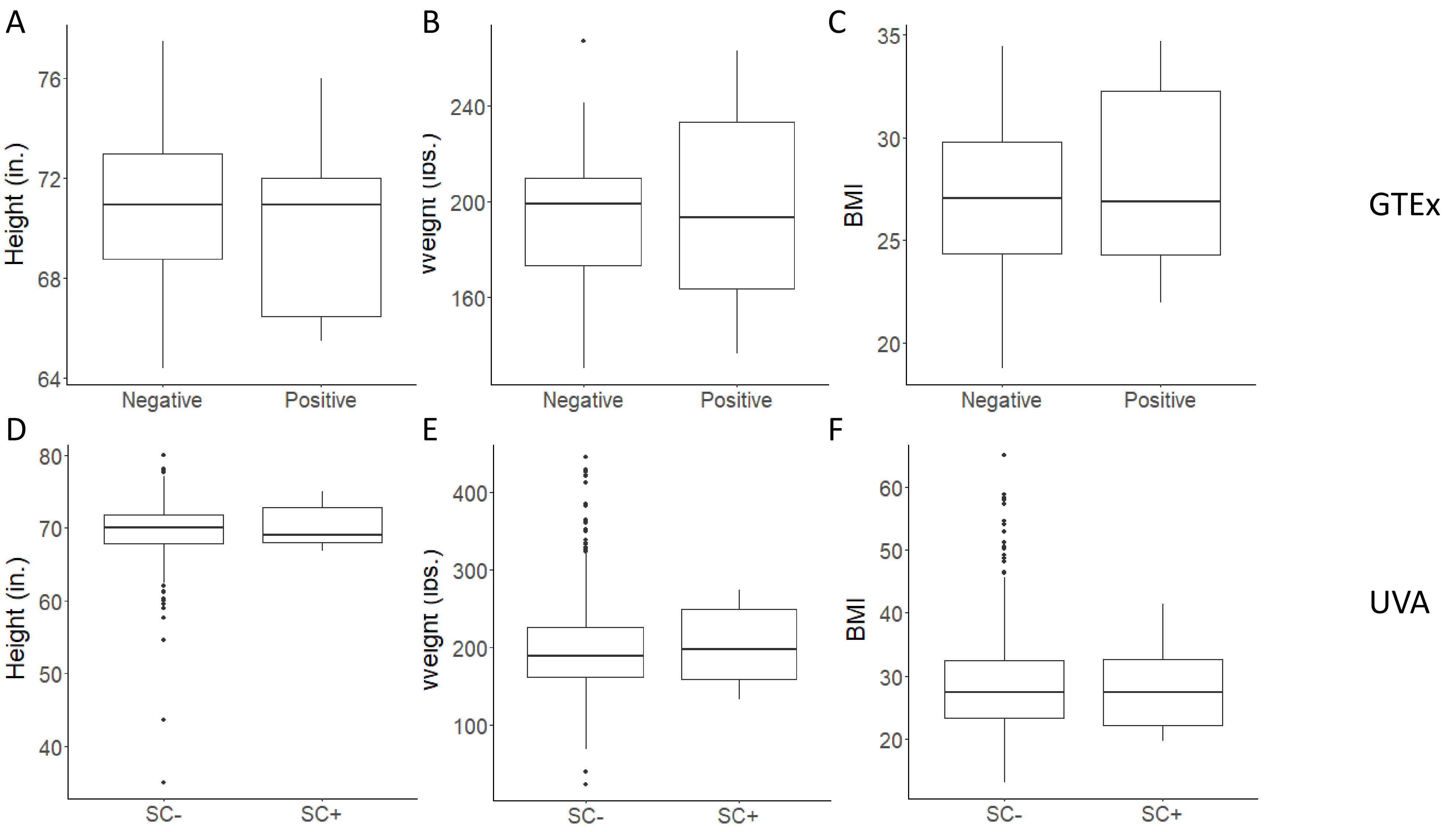

Supplementary Figure S4

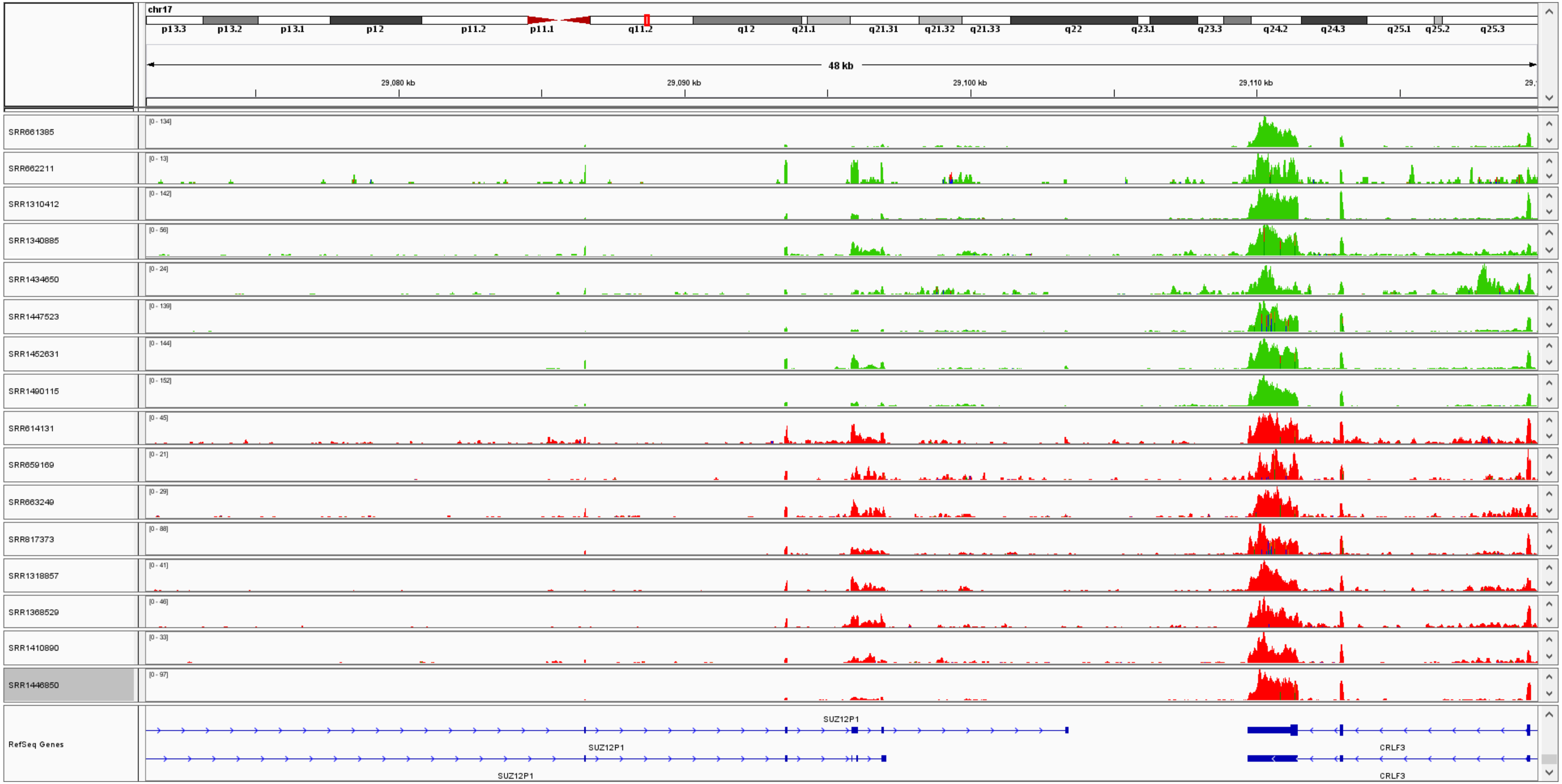

Supplementary Figure S5

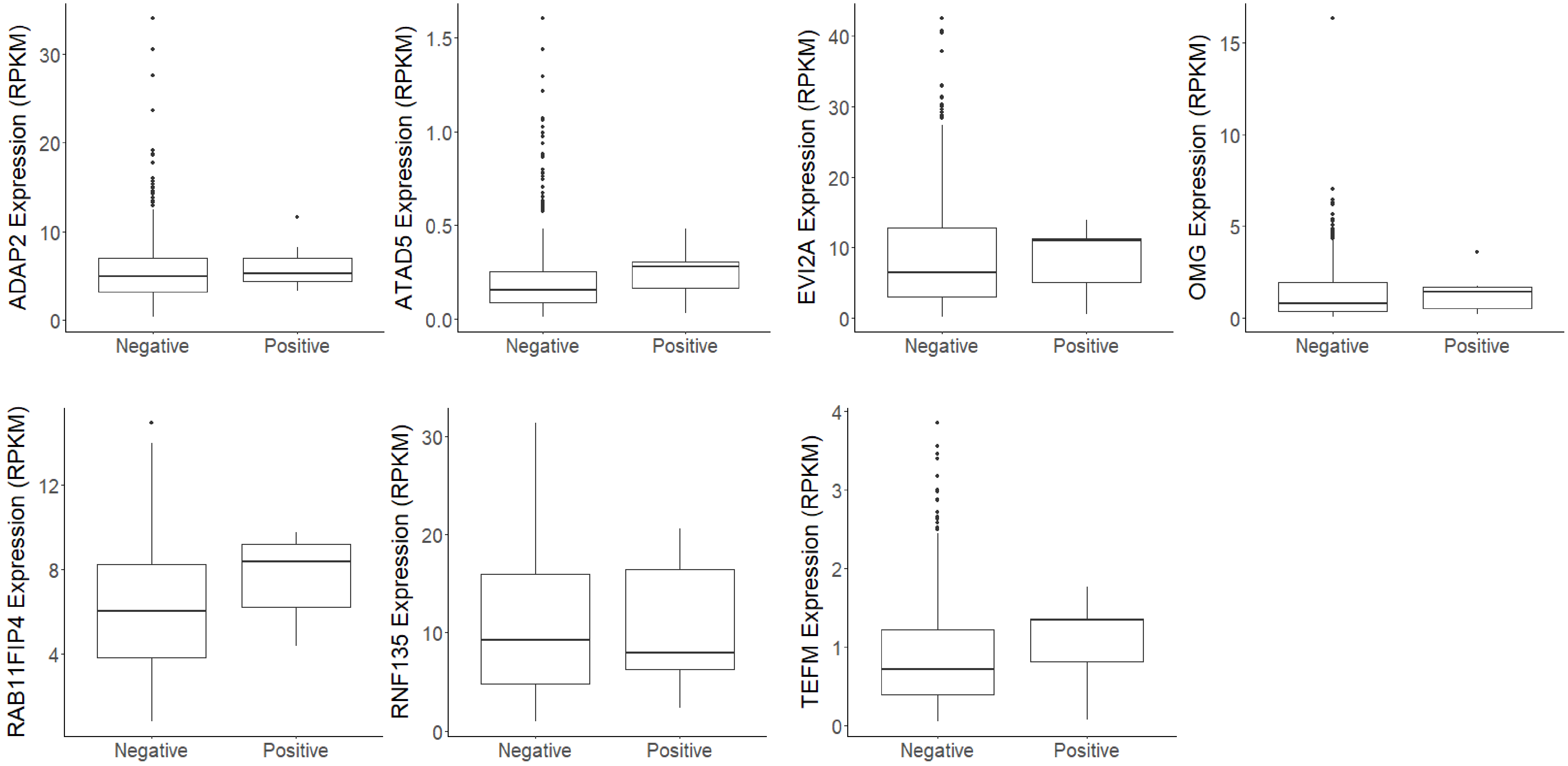

Supplementary Figure S6

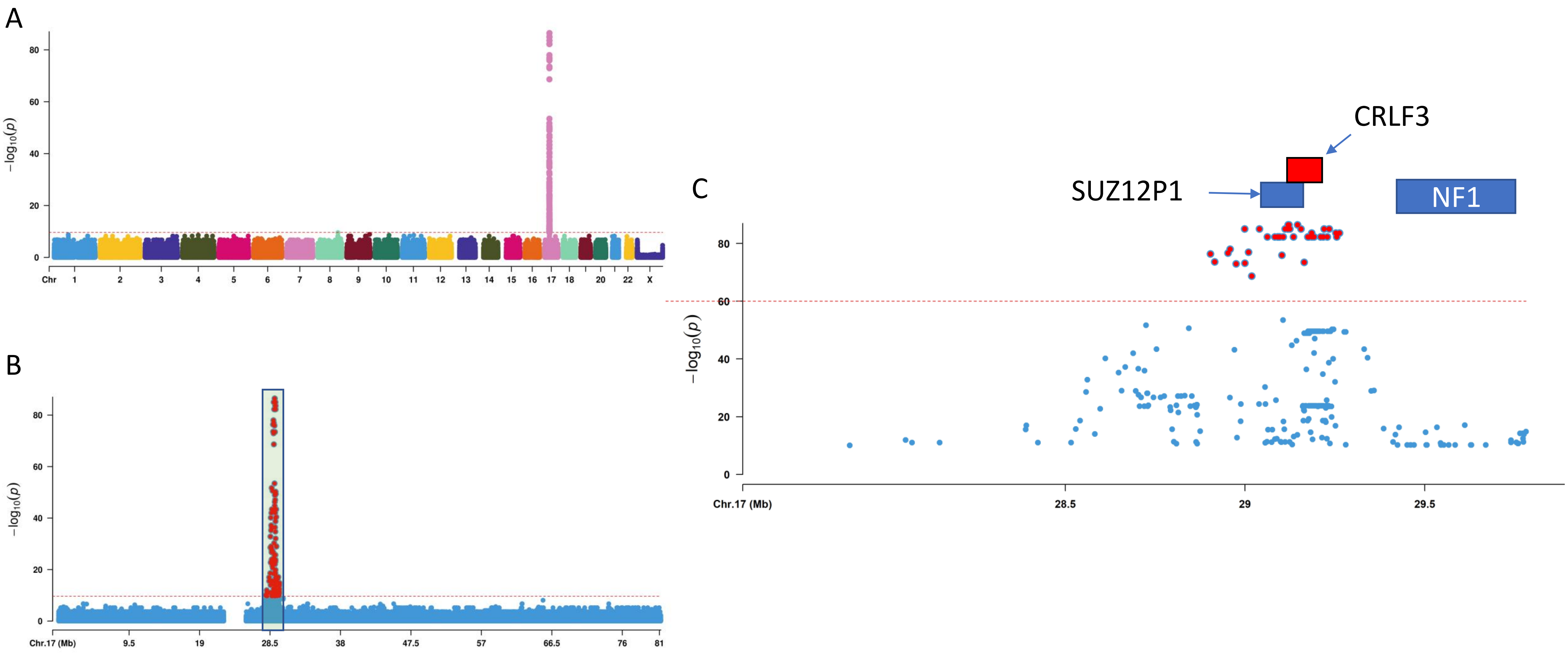

Supplementary Figure S7

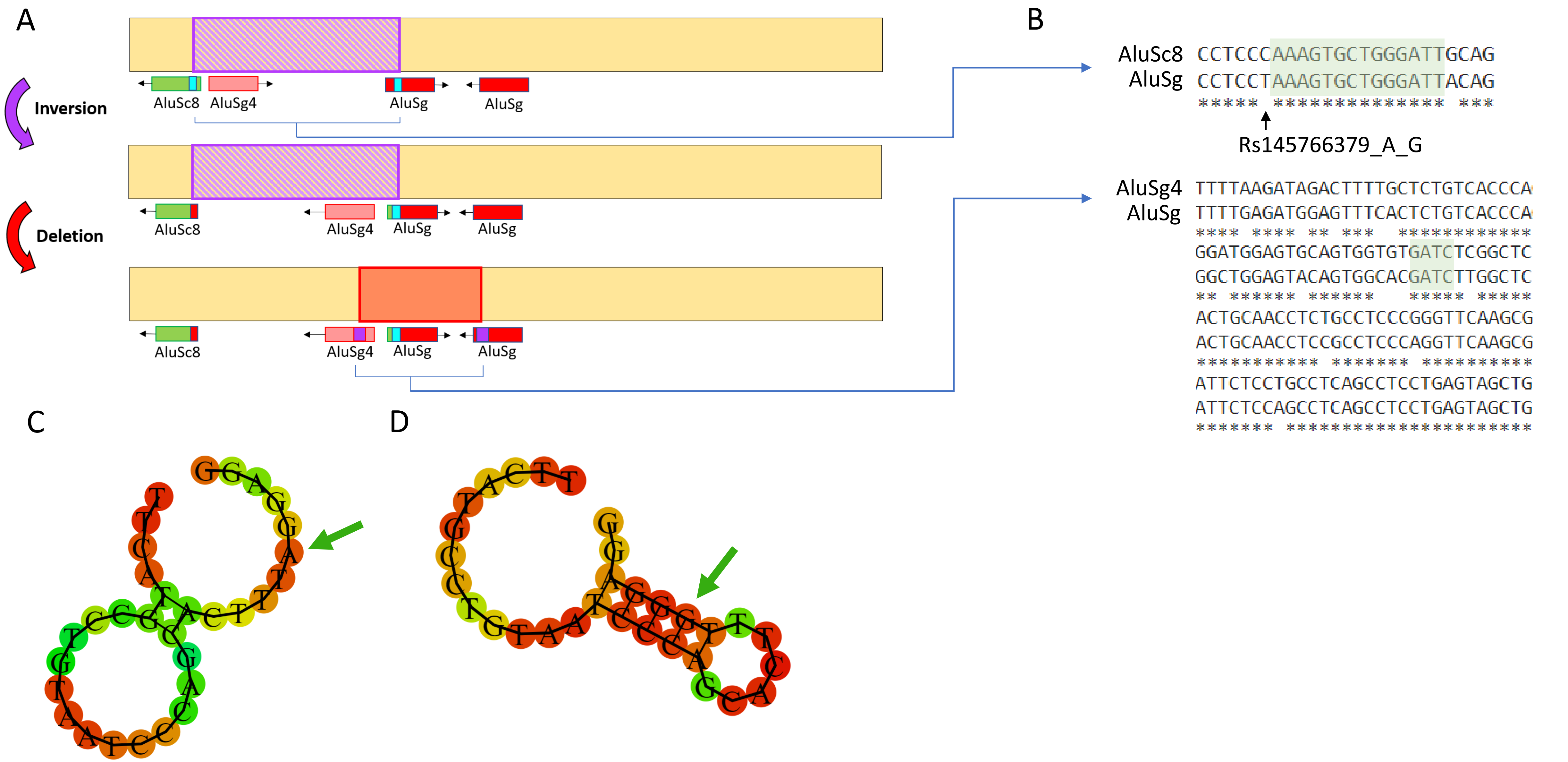
